# Feathers of Grace: The “After You” Gesture in Japanese Tits

**DOI:** 10.1101/2024.04.29.591630

**Authors:** Sergio Da Silva, Raul Matsushita

## Abstract

A recent study found evidence of symbolic gesture use in Japanese tits (*Parus minor*). The study reveals how these birds use wing-fluttering movements to transmit an “after you” to their partners, implying a degree of cognitive skill previously thought to be unique to humans and great apes. If confirmed, this research contradicts long-held notions about animal communication by proving that Japanese tits not only participate in intricate vocal communications, which can comprise phrases with specific grammatical rules, but also use body language as a form of engagement. Here, we evaluate this claim by inspecting the data and utilizing bootstrapping to expand the sample size. We find a large variation in how frequently the “after you” gesture is employed in different bootstrap samples, suggesting that while the gesture is a consistent behavior, its frequency can vary significantly. Moreover, the timing of the male’s response to the female’s gesture can fluctuate, though the response itself appears to be a stable phenomenon. Because the frequency of the “after you” gesture varied greatly, especially in light of potential cognitive and other biases influencing the study, this bold claim should be taken with caution.

## 1. Introduction

A study just reported in *Current Biology* [1] presents evidence of symbolic gesture use among the Japanese tit (*Parus minor*). The research reveals how these birds, through wing-fluttering gestures, communicate an “after you” to their mates, suggesting a level of cognitive skill previously believed to be exclusive to humans and great apes [2−4]. If robust, this discovery challenges conventional assumptions about animal communication, suggesting that Japanese tits not only engage in elaborated vocal communications, which can include phrases with particular grammatical rules, but also utilize body language as a form of interaction. The aim of this communication is to assess this claim.

The researchers observed eight breeding pairs of Japanese tits over 321 nest visitations, noting that the gesture of wing fluttering was primarily used by females to signal to their males to enter the nest box first, even if the female arrived first. This behavior was found to be symbolic [5] because it was specifically aimed at the mate rather than the nest box, was only observed in the presence of the mate, and ceased once the mate entered the nest, indicating a sophisticated form of communication beyond mere pointing or other deictic gestures [5, 6]. “After you” is a symbolic gesture, which is more cognitively demanding then deictic gestures [5]. Deictic gestures in animals are communicative movements or signals that specifically direct the attention of others to objects or events in the environment.

The implications of the study are deep, extending beyond ornithology to the broader fields of animal cognition and communication. By showing that birds are capable of symbolic gestures, the research not only potentially improves our understanding of avian intelligence but also prompts a reevaluation of the evolutionary origins of communication and language. The hypothesis that bipedalism in humans led to the development of gestures, as suggested by the authors, draws a parallel to birds, whose freed wings while perching may facilitate gestural communication. The connection between human and avian gestural communication underscores the potential for further discoveries in animal behavior, offering insights into the cognitive abilities of species beyond our own and improving our understanding of the evolution of sophisticated communication systems, including language.

We begin evaluating the images and videos provided in the report. The report’s images and videos suggest a bird making a possible “after you” gesture, although it is unclear. One photo captures a bird with wings spread, which could indicate gesturing, landing, or taking off. Another image shows a bird perched, possibly indicating feeding, signaling, or balancing. Possible interpretations include: (1) Feeding gestures, hinting at food presence or sharing rituals. (2) Mating displays, with wing fluttering for courtship. (3) Aggressive displays, using wings for warnings or territorial claims. (4) Balancing movements for stability against wind or during landings. In their Figure 1B, the authors addressed how social contexts and sex influence nest visitations. They noted that in the “with a mate” context, the frequency of visits differed between males and females, with instances where one parent made multiple visits while the other remained outside the nest box with food. However, the images and videos provided are inconclusive.

**Figure 1.**
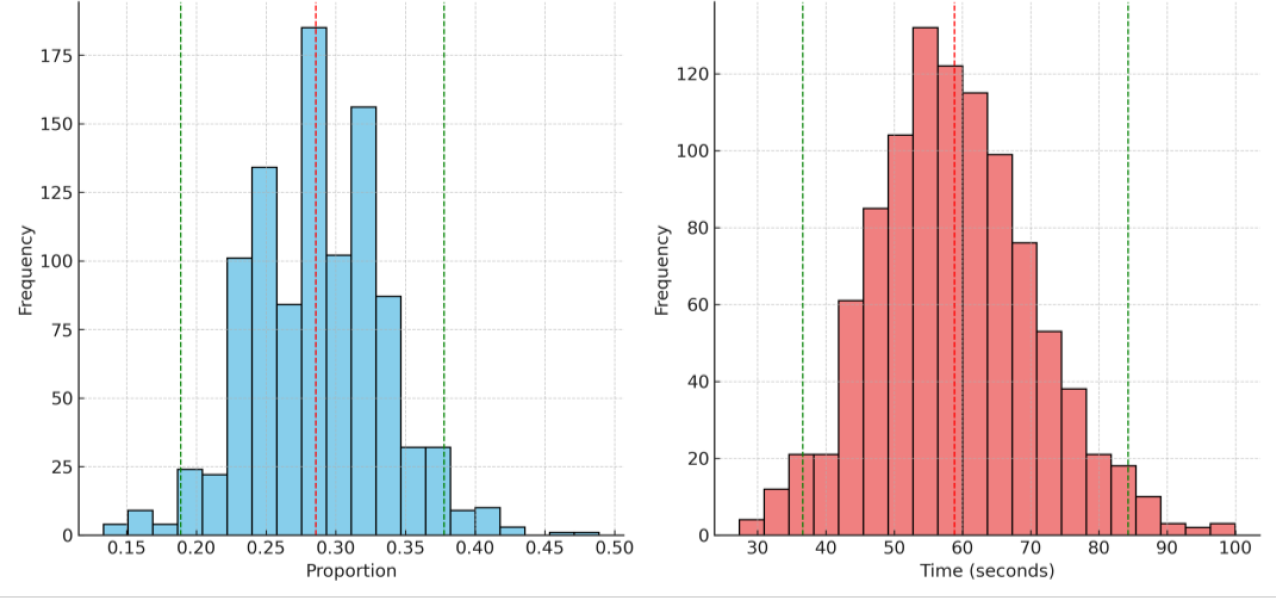
Left panel: shows the distribution of the proportion of the “after you” gesture when mates are present. The dashed red line indicates the mean proportion across the bootstrap samples, while the green lines represent the 95% confidence interval. Right panel: displays the distribution of mean latency times for males to enter the nest after the gesture. Similar to the left panel, the dashed red line shows the mean latency, and the green lines mark the 95% confidence interval.

Furthermore, the narrative of one video contained within the supplementary material appears to be susceptible to the influence of a narrative fallacy [7], making the “after you” gesture’s certainty challenging to confirm without specific context and actions analysis. First, it is unclear if the wing-fluttering is exclusive to the mate’s presence and directed solely at them, rather than the surroundings or nest box. Second, the video implies the female continues her gesture even after the mate enters the nest, questioning the gesture’s cessation and purpose. Lastly, the video does not clarify whether this gesture is a consistent behavior pattern or an isolated incident.

Another issue is the risk of anthropomorphizing [8, 9]. This is a valid concern in animal behavior studies. Anthropomorphizing occurs when human traits, emotions, or intentions are attributed to nonhuman entities, often leading to misinterpretation of animal behavior. We think the authors took measures to avoid anthropomorphizing and support their claim. When observing the 321 nest visits, they paid special attention to circumstances in which the birds encountered their mates near the nest. They found that wing-fluttering behavior occurred significantly more often when a mate was present as opposed to when the bird was alone, and that this behavior prompted the mate to enter the nest first, without any direct physical contact. The authors were careful to define gestures according to established criteria in animal behavior studies, ensuring that the gestures were goal-directed, did not act as direct physical agents, and elicited a specific response. They analyzed the effect of the wing-fluttering on the order of nest entry and the latency of the males to enter after the gesture was observed, using statistical models to assess the significance of their finding. By observing that the wing-fluttering ceased after the mate had entered the nest and was always directed towards the mate (not the nest entrance itself), they argued that this behavior fulfills the criteria of a symbolic gesture rather than a deictic one.

We argue that the study’s methodology, while appropriate, has inherent limitations and potential cognitive and other biases. Here, we detect ten of these: (1) The “law of small numbers” [10]. This refers to sample size and generalizability: Observations from 321 nest visits involving eight pairs of birds may not represent all Japanese tits or other species, limiting the finding’s applicability. (2) Observer expectancy effect [11]: Researchers’ expectations might unconsciously bias which behaviors are noted, possibly overlooking other significant behaviors due to confirmation bias [12]. (3) Anthropocentric interpretation [8, 9]: Efforts to avoid humanizing animal behavior can still be biased by human perspectives on gesture and communication. (4) Contextual variability [13]: Wing-fluttering observed during nest visits might have different meanings in other contexts, questioning the gesture’s universality. (5) Ecological validity [14, 15]: Using nest boxes for observation might not accurately reflect natural environments, potentially altering bird behavior. (6) Cultural transmission bias [16]: If gestures are learned, unnoticed cultural variations within Japanese tits could affect the gesture’s prevalence and meaning. (7) Experimental constraints [17, 18]: Study conditions might impose unnatural behaviors on the birds, influencing outcomes. (8) Statistical artifacts [19, 20]: Significant results could stem from the analysis methods rather than actual biological phenomena, particularly with multiple comparisons. (9) Publication bias [21]: A tendency to publish positive findings over negative or inconclusive ones could misrepresent the frequency of elaborated gestural communications among birds. (10) Temporal and geographic constraints [22]: The study’s specific time and location limit understanding of how behaviors vary with environmental and social changes.

To overcome these limitations, it is of course recommended to conduct further studies on larger and varied populations, in diverse settings, and possibly through longitudinal research to validate and broaden the finding. Additionally, peer reviews, independent replications, and meta-analyses are also important for confirming these results and addressing potential cognitive and other biases. The authors themselves admit that further studies are necessary.

The authors employed a Generalized Linear Mixed Model (GLMM) [23] and a Cox mixed-effects model [24] to analyze their data. These statistical models are designed to handle the complexities of repeated measures on the same subjects (the birds), which is common in behavioral studies. They reported *p*-values for the different tests they conducted. For instance, the presence of wing-fluttering when a mate was nearby had a *p*-value of less than 0.0001, indicating that the results were highly unlikely to have occurred by chance. They also analyzed the effect of wing-fluttering on the order of nest entry (*p* = 0.0005) and the latency of males to enter the nest (*p* = 0.0035), both of which were statistically significant. These methods indicate that the authors considered the possibility that their results occurred by chance and employed an appropriate statistical analysis to minimize this likelihood. The low *p*-values suggest that the observed behaviors were not likely due to chance, giving confidence in the robustness of their finding. Nevertheless, continued replication of the study by other researchers would add to the validity of the finding by confirming whether the results can be consistently observed across different samples and conditions.

The supplemental material of the paper outlines the experimental methods but also calls attention to potential study limitations and areas for improvement, including: (1) Temporal and weather restrictions: The observations were made during specific times and under specific weather conditions, which may not capture the full range of natural behaviors exhibited by the birds. Future studies could aim to observe the birds under a wider range of conditions. (2) Behavioral classification validity: While the classification of wing-fluttering behavior was confirmed with a perfect agreement among hypothesis-blind observers, it is still a human interpretation of animal behavior, which may not fully capture the birds’ intentions. (3) Observer influence: the authors maintained a 15-meter distance; however, despite these precautions, human presence could still unpredictably affect animal behavior. Though the researchers confirmed that the birds made at least one feeding visit, there is still a potential for altered behavior due to human observation. (4) Sample size: The study used eight pairs of birds for observations, which is a relatively small sample size and may not capture the full diversity of behaviors across the species. (5) Statistical considerations: They have taken care to use appropriate statistical methods, including Firth’s [25] bias reduction for maximum likelihood estimates due to complete separation, and checking for multicollinearity with variance inflation factors. However, any statistical analysis is based on the assumption that the model correctly represents the underlying biological processes.

The supplemental material also hints at underlying cognitive biases, such as researchers possibly favoring data that supports their hypotheses and the selection of nest-box-using birds not fully representing the wild population. Additionally, interpreting wing-fluttering as a polite gesture might be projecting human concepts onto animal behavior. To overcome these limitations, future research could benefit from increasing the number of bird pairs studied to better represent the species and extending the observation period to capture evolving communication patterns. Employing observers unaware of the study’s hypotheses could ensure unbiased behavior classification. Comparing the gesture with those of other bird species could help determine its specificity or universality. Experimentally altering conditions, like the presence of a mate, could offer insights into the gesture’s intent. Finally, conducting studies in various environments and seasons would assess the behavior’s consistency across different contexts, providing a more comprehensive understanding of the phenomenon.

To assess the authors’ claim, we examine their data. We look at synthetic ways for expanding their data sample. Increasing the sample size in a synthetic way typically involves data augmentation, a technique often used in machine learning to artificially expand the dataset. In the context of behavioral studies, however, creating synthetic observations that are valid and reliable enough to augment actual behavioral data is challenging, as it requires not only duplicating data but simulating new, realistic behaviors that are consistent with observed patterns. Several hypothetical methods for data enlargement come with their own set of significant caveats. Bootstrapping [26] involves resampling existing data to create new datasets for estimating variability and bolstering statistical inferences, though it falls short in generating genuinely new data or diversifying behavioral patterns. Agent-based modeling [27] offers a creative solution by employing AI to simulate bird behaviors like nest visitation and wing-fluttering, potentially creating realistic, novel data points for analysis. Monte Carlo simulations [28], leveraging known probabilities of various behaviors, use random generation based on probability distributions to simulate additional datapoints. Techniques of interpolation and extrapolation [29] apply statistical or machine learning methods to generate datapoints within or slightly beyond the observed behavioral spectrum, drawing on existing patterns. Lastly, the Synthetic Minority Over-sampling Technique (SMOTE) [30], designed for balancing datasets in machine learning, could be adapted to generate conceivable new behaviors by interpolating between observed instances, providing a fresh perspective on behavioral data.

Synthetic data augmentation in behavioral studies is not commonly practiced because behaviors are complex and context-dependent, making it difficult to generate synthetic behaviors that are indistinguishable from real ones. Standards in animal behavior research usually require observations of actual behavior, as synthetic data may not capture the full complexity of animal interactions. Furthermore, the unpredictability and variability of animal behavior make it hard to create realistic synthetic data without introducing biases or artifacts that could skew results. Therefore, the best approach to increase the sample size and the robustness of the findings is usually to conduct additional observations, either by expanding the study to include more pairs or by replicating the study in different locations or populations. If synthetic methods are to be used, they should be carefully validated against real-world data to ensure they are producing realistic and valid behaviors. Having said that, we chose the bootstrapping strategy. It is the most applicable and conservative approach to the data at hand. It can help estimate the variability and confidence intervals of the observed behaviors without assuming behaviors that were not observed. It does not create new behaviors, but it can provide insights into the stability and robustness of the results obtained from the sample. For example, it could be used to assess the robustness of the finding that the “after you” gesture is significantly associated with the mate’s subsequent behavior.

## 2. Materials and Methods

We considered the authors’ provided data at https://data.mendeley.com/datasets/256z7k654k/1 along with the steps for reproduction suggested by them.

Considering the datasets (wing-fluttering behavior and nest entry times) for analysis, we created multiple bootstrap samples from the original data. Each sample was of the same size as the original dataset but were drawn with replacement. This means some observations may appear more than once, while others may not appear at all in the bootstrap sample. Thus, we created numerous bootstrap samples from the original data. Each sample randomly selected observations with replacement, simulating the effect of drawing from the original population.

For each bootstrap sample, we computed the frequency of wing fluttering when mates are present and the latency times for males to enter the nest after the females’ gesture. Therefore, for each bootstrap sample, we calculated the mean latency for males to enter the nest after the “after you” gesture and the proportion of times this gesture occurs when the mate is present. Then, we calculated the confidence intervals for these statistics across all bootstrap samples. This provides insight into their variability and how they might differ by chance alone. We calculated the 95% confidence intervals for these statistics from the bootstrap samples to assess their stability. Next, we interpreted the results to see if the observed behaviors in the original dataset (like the significant association of the gesture with subsequent behavior) hold up under the resampling, which would suggest a robust finding.

## 3. Results

A 95% confidence interval means if we were to take many samples and calculate a confidence interval for each, approximately 95% of these intervals would contain the true population parameter. The bootstrapping analysis provided the following 95% confidence intervals for our key statistics: (1) Proportion of “after you” gesture when mate is present. We found a confidence interval of [18.9%, 37.8%]. This indicates that in bootstrap samples, the proportion of observed “after you” gestures (when mates are present) varies from about 18.9% to 37.8%. Therefore, there is large variation in how frequently this gesture is employed in different bootstrap samples, suggesting that while the gesture is a consistent behavior, its frequency can vary significantly. (2) Mean latency for males to enter the nest. We found a confidence interval of [36.6, 84.3] seconds. This means that the average time males take to enter the nest after encountering the female varies between 36.6 and 84.3 seconds in the bootstrap samples. Therefore, this variability emphasizes that the timing of the male’s response to the gesture can fluctuate, though the response itself appears to be a stable phenomenon. Figure 1 provides a clear visualization of the variability and the central tendencies in the data, helping to illustrate the conclusions derived from the bootstrapping analysis.

Our previous analysis ignores one piece of additional information: wing-fluttering behavior is context-dependent. When the female is the first feeder, wing fluttering occurs about 3.03% of the time. In contrast, when the female is not the first feeder, the occurrence of wing fluttering jumps to approximately 95.83%. Fisher’s exact test *p*-value of 10^−13^ from these proportions indicates a significant difference based on whether the female is the first feeder or not, suggesting a strong relationship between wing fluttering and feeding order among females when the mate is present.

The cross table between wing-fluttering behavior vs. sex gives a Fisher’s exact test *p*value of 0.0002, and the bootstrap analysis indicates that the average proportion of female birds displaying wing-fluttering behavior when a mate is present is approximately 42.09%. The 95% confidence interval for this proportion ranges from 29.82% to 54.39%. Therefore, the behavior varies greatly amongst bootstrap samples. The dashed lines in Figure 2 represent the 95% confidence interval, and the solid blue line shows the mean proportion.

**Figure 2.**
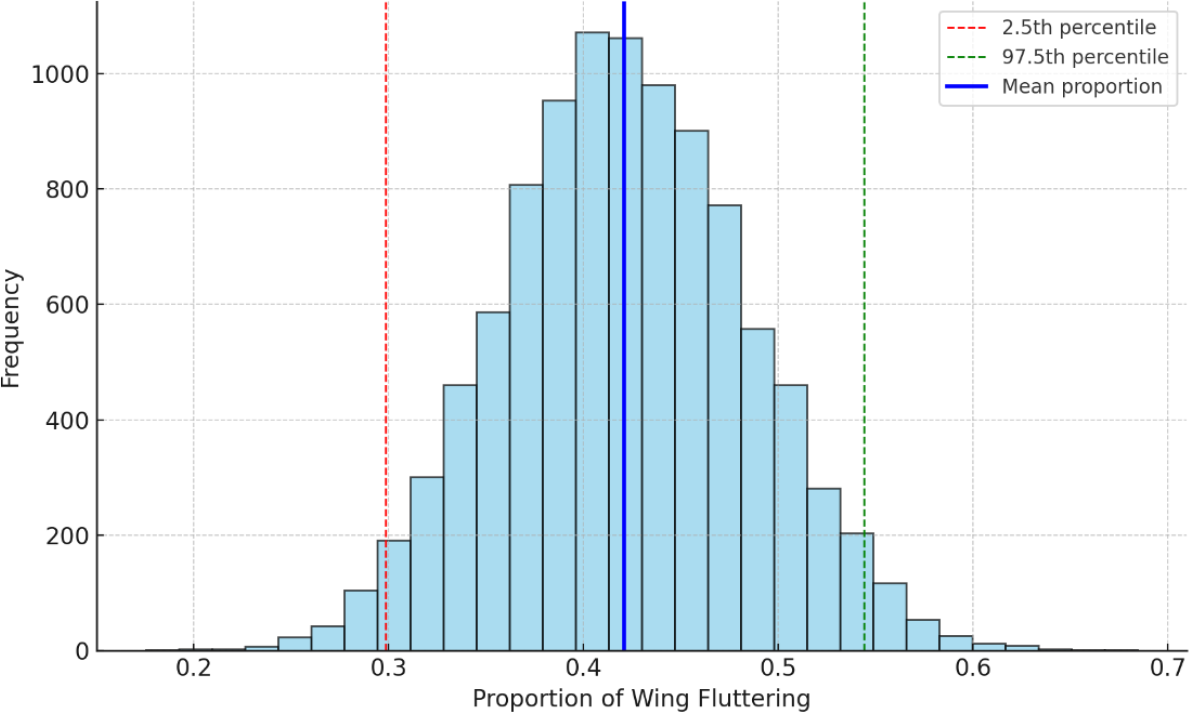
Bootstrap distribution of wing-fluttering proportions for females. This figure visualizes the distribution of the bootstrap samples for the proportion of female birds engaging in wing-fluttering behavior when a mate is present. Behavior in bootstrap samples varies significantly.

## 4. Discussion

Now we evaluate the implications of our results in the context of the paper’s claim.

Regarding statistical significance, the authors’ work should have supplied statistical tests (such as nonparametric tools like Fisher’s exact test and the bootstrap method) comparing scenarios where the gesture is used vs. not used, to robustly claim the significance of these behaviors’ effects. However, the low *p*-values reported in the paper suggest that these differences are statistically significant, adding weight to the claim.

They should also have evaluated effect size and practical significance. Understanding how much quicker males enter the nest when the gesture is made, compared to when it is not, is very important. A significant but small difference might not be biologically meaningful, despite its statistical significance.

Another factor neglected by the authors was the consistency across samples. Our bootstrapping results showing some degree of variability in how the gesture impacts male behavior (seen in the range of latency times) suggest that while the effect is generally consistent, environmental or contextual factors could influence the strength and timing of the response.

Finally, the variation noted in the bootstrap analysis emphasizes the need for cautious interpretation and further studies to explore this behavior under different environmental conditions or in different populations to affirm its generality and significance.

## 5. Conclusion

The authors’ suggestion that Japanese tits utilize gestures to say “after you” to their mates by fluttering their wings is a bold claim with far-reaching cognitive and evolutionary ramifications. If the females’ “after you” gesture consistently has a significant influence on male behavior, it may indicate a more complicated level of communication and cognitive abilities than is normally assumed in birds. It indicates a link with symbolic gestures in apes, implying evolutionary convergence in communication mechanisms across vastly divergent lineages.

We discussed a variety of cognitive and other biases that could be affecting such a claim. Because cognitive biases cannot be corrected after the completion of the study, we focused on the “law of small numbers” bias, which refers to jumping to conclusions based on small samples. Because the authors’ sample is too small, we opted for a bootstrapping strategy to expand their datasets and reevaluate their claim. The bootstrapping analysis could not disconfirm the paper’s claim that the “after you” gesture has significant effects on male behavior, indicating its potential as a symbolic gesture rather than a simple reflexive or situational response. However, the observed variation we also found suggests a more nuanced interpretation and emphasizes the significance of future study to validate and investigate the depth and consistency of this intriguing behavioral feature.

## Data Availability Statement

The data are available at https://data.mendeley.com/datasets/256z7k654k/1. Accessed on 14 April 2024.

## Appendix

Does the presence of wing fluttering behavior in female Japanese tits cause a reduction in the time it takes for their mates (males) to enter the nest? To answer this, we employ causal inference [31, 32] and formulate a hypothesis:

### Hypothesis

Wing fluttering by females leads to a quicker response by males, entering the nest more swiftly, indicating a form of effective communication.

The data involves observations of Japanese tits, focusing on their wing-fluttering behavior as a form of communication, particularly signaling “after you” to their mates. The two datasets provided by the authors track: (1) Wing fluttering behavior: Records whether wing fluttering occurred, the sex of the bird, and whether the mate was present. (2) Nest entry time: Tracks the time it takes for the males to enter the nest after the wing fluttering gesture by the females.

We define variables as follows: (1) Exposure variable: Presence of wing fluttering behavior (binary: yes/no). (2) Outcome variable: Time for male Japanese tits to enter the nest after the gesture (continuous: measured in seconds).

Potential confounders in the analysis could include the time of day, as behavioral patterns may vary throughout the day, affecting both the observed behavior and the resulting actions. The physical characteristics of the nest location could also influence the likelihood of wing fluttering and the speed of the male’s response. Additionally, the presence of a mate at the time of the gesture could affect the behavior of both the female making the gesture and the male’s response time.

The wing fluttering data includes observations on various dates, where each entry documents an observation number, the order in which birds arrived at the nest, and the sex of the bird. It also notes whether wing fluttering occurred, if a mate was present during this behavior, and identifies which bird was the first to feed if applicable. Additionally, each record is associated with a specific nest and bird through unique identifiers.

The nest entry time data records observations that include the date, details about each bird and nest through unique identifiers, and the observation number. It also notes the order in which birds arrived at the nest, whether wing fluttering was observed during this particular event, and the time it took for the male to enter the nest following a female’s gesture, measured in seconds. The status of each observation is also noted, although its specific definition is not provided in the data sample.

The next steps in data preparation involved merging the datasets based on common attributes to align occurrences of wing fluttering with corresponding response times. The data was cleaned by filtering out irrelevant observations and managing any missing entries. Following this, a statistical analysis was conducted to explore the relationship between the presence of wing fluttering and the males’ response time. This process prepared the data for further analysis.

After merging and cleaning the data, there were 33 records available for analysis. The majority of these observations, about 94%, indicated that there was no wing fluttering, with only 6% showing occurrences of wing fluttering. The time it takes for males to respond, known as the time to male feed, had a mean of approximately 59 seconds and a standard deviation of about 68 seconds, highlighting a large variability in response times, which range from 7 to 180 seconds.

To explore the causal relationship, we performed a statistical analysis comparing the response times between instances with and without wing fluttering. This helped determine if the presence of wing fluttering significantly affects the time it takes for males to enter the nest.

The *t*-test comparing the response times between instances with wing fluttering and those without yielded the following results: *t*-statistic: −3.30; *p*-value: 0.0048. This *p*-value is less than the typical significance level of 0.05, indicating that the differences in response times between the two groups are statistically significant. Therefore, we can reject the null hypothesis that there is no difference in mean response times between the groups, suggesting that wing fluttering does have an effect on the time it takes for males to enter the nest.

The histogram in Figure A1 shows distinct distributions for the two groups. The majority of responses without wing fluttering (blue) have a wider range and are generally higher, indicating longer response times. Responses with wing fluttering (red) are fewer (given the smaller number of observations) and are concentrated at lower times, indicating quicker responses.

**Figure A1.**
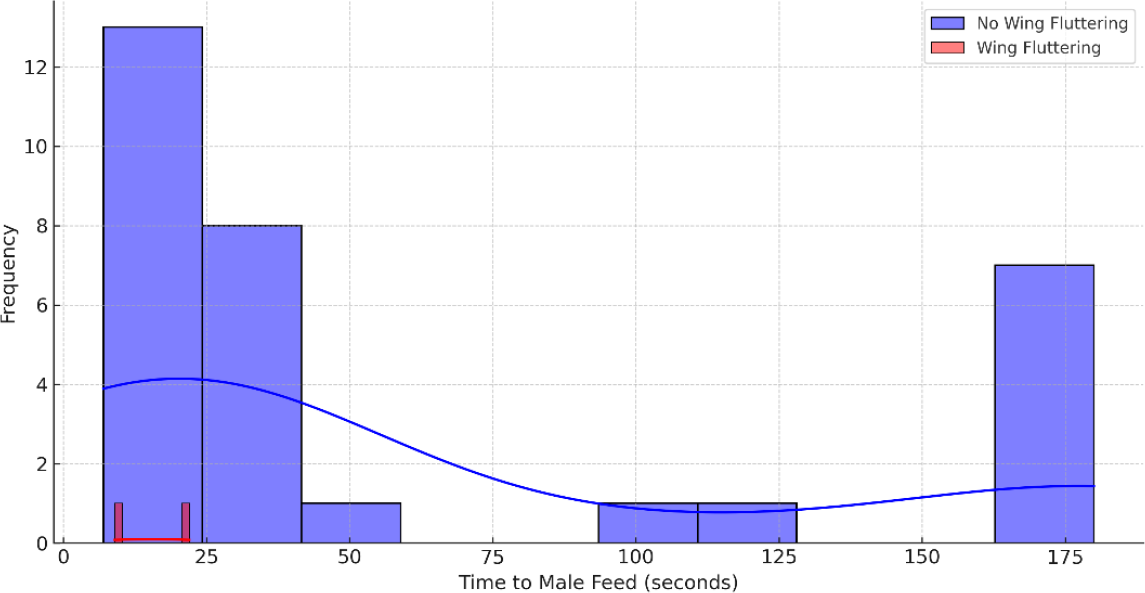
Distribution of response times with and without wing fluttering.

As a result, the analysis supports the hypothesis that wing fluttering by female Japanese tits tends to lead to quicker responses from their male mates, entering the nest more swiftly. This suggests that wing fluttering could be an effective communicative gesture in this species.

This analysis provides statistical evidence suggesting that the “after you” gesture by female Japanese tits (indicated by wing fluttering) does indeed result in a statistically significant reduction in the time it takes for males to enter the nest. This can be interpreted as the males responding more quickly when this gesture is made. However, a few considerations are necessary before concluding robustness. First, the number of instances with wing fluttering was relatively small compared to those without. A larger sample size, especially more observations with wing fluttering, would help to confirm these findings. Second, although the analysis accounted for the presence of wing fluttering, other factors could influence response times. Further analysis would be needed to adjust for potential confounders like time of day, environmental conditions, and individual differences between birds. Third, robust findings should ideally be reproducible across different studies or similar analyses with new data sets. Confirming these results in different contexts or with additional data would strengthen the claim of robustness. Lastly, beyond statistical significance, understanding the ecological or biological implications of these gestures in the natural behavior and life cycle of Japanese tits is crucial for interpreting the robustness of the “after you” gesture’s significance.

In conclusion, while the current analysis provides good preliminary evidence supporting the potential communicative value of the “after you” gesture, further studies with larger and more diverse samples, consideration of confounders, and replication of results are recommended to firmly establish the robustness of this finding.

